# Quantitative assessment of association between noncoding variants and transcription factor binding

**DOI:** 10.1101/2022.11.22.517559

**Authors:** Ningxin Ouyang, Alan P. Boyle

## Abstract

Association fine-mapping of molecular traits is an essential method for understanding the impact of genetic variation. Sequencing-based assays, including RNA-seq, DNase-seq and ChIP-seq, have been widely used to measure different cellular traits and enabled genome-wide mapping of quantitative trait loci (QTLs). The disruption of cis-regulatory sequence, often occurring through variation within transcription factor binding motifs, has been strongly associated with gene dysregulation and human disease. We recently developed a computational method, TRACE, for transcription factor binding footprint prediction. TRACE integrates chromatin accessibility and transcription factor binding motifs to produce quantitative scores that describe the binding affinity of a TF for a specific TFBS locus. Here we have extended this method to incorporate variant data for 57 Yoruban individuals. Using genome-wide chromatin-accessibility data and human TF binding motifs, we have generated precise, genome-wide predictions of individual-specific transcription factor binding footprints. Subsequent association mapping between these footprints and nearby regulatory variants yielded numerous footprint-variant pairs with significant evidence for correlation, which we call footprint-QTLs (fpQTLs). fpQTLs appear to affect TF binding in a distance-dependent manner and share significant overlap with known dsQTLs and eQTLs. fpQTLs provide a rich resource for the study of regulatory variants, both within and outside known TFBSs, leading to improved functional interpretation of noncoding variation.

## Introduction

Transcription factors (TFs) recognize and bind to specific DNA sequences within cis-regulatory elements, known as transcription factor binding sites (TFBSs), to perform their regulatory activities. Although TFBSs make up only 8% of the genome, they contain 31% of genome-wide association study (GWAS) variants, suggesting they play a largely underappreciated role in human disease (ENCODE Project Consortium 2012). Binding of TFs to TFBSs is fundamental for transcription regulation and the effect of regulatory variants on these regions is directly interpretable. The functional effect of noncoding single nucleotide variants (SNVs) is often observed as a strengthening or weakening of individual TF binding activity, thus identifying the effects of sequence variations on TF binding is key to understanding and interpreting the downstream consequences on gene expression and disease phenotypes.

Likewise, association mapping of quantitatively measurable molecular traits is a powerful approach for studying genetic variation. Genome-wide mapping of expression quantitative trait loci (eQTLs) examine the impact of variants on gene expression levels (Cheung et al. 2005; Veyrieras et al. 2008), identifying variants with potential gene-regulatory impact. However, because variants in close proximity are often in linkage disequilibrium, it is difficult to determine which variant(s) are directly responsible for differential gene expression, let alone discern the underlying regulatory mechanisms driving these associations. One putative mechanism is that variants can alter the likelihood and/or strength of TF binding to an individual TFBS, thereby affecting regulatory networks. Previous studies have demonstrated that mutations in TFBSs alter TF binding affinity, and are associated with human diseases including cancer and type 2 diabetes (Gaulton et al. 2010).

Binding of TFs to TFBSs often protects underlying DNA from DNaseI digestion, leaving behind a distinct digestion pattern known as a footprint. DNase I hypersensitive site sequencing (DNase-seq) measures open chromatin regions where DNase I cuts at higher frequencies, allowing for genome-wide footprinting. We have recently developed a computational footprinting method, TRACE that simultaneously annotates the genome with predicted footprints for multiple TFs (Ouyang and Boyle 2020). The TRACE model integrates both DNA accessibility (DNase-seq or ATAC-seq) and genome sequence information (PWMs to predict which TF binds each predicted footprint), into a quantitative binding score that can be compared across individual genomes. Thus, changes in binding activity caused by variation in genotype and/or chromatin accessibility between individuals will manifest as differences in the TRACE score at a given position. Therefore, TRACE scores can be used as a quantitative trait to identify variants associated with changes in TF binding strength between individuals, which we term “footprint QTLs”, or fpQTLs.

Here we leverage DNase-seq, RNA-seq, and HapMap variant data from 57 Yoruban lymphoblastoid cell lines (LCLs) to enable genome-wide fpQTL identification and examine their impact on phenotypic variation (1000 Genomes Project Consortium et al. 2010; International HapMap Consortium et al. 2007; Pickrell et al. 2010; Degner et al. 2012). This genome-wide mapping of fpQTLs provides additional information for the functional interpretation of human noncoding variants, thus enabling further insights into the mechanisms underlying population-level regulatory variation.

## Results

### Genome-wide identification of fpQTLs

Genetic variation can alter TF binding sequences and therefore affect the likelihood and/or strength of TF binding. As an example, the alternative allele for rs1338681, is known to disrupt a CTCF binding motif (**Figure 1C**), leading to reduced binding affinity (**Figure 1D**). However, it is also possible for variants outside a footprint to directly influence TF binding at nearby TFBSs, e.g., by altering local chromatin accessibility. Identifying these variants requires a quantitative score capable of expressing a TFs likelihood of binding at any given footprint. Our recently-published TRACE method employs chromatin accessibility data and TFBS motif models to generate quantitative scores summarizing the likelihood of TF binding at any given footprint. These scores can be compared across individuals in order to evaluate the impact of nearby variants on TF binding activity.

**Figure 1:**
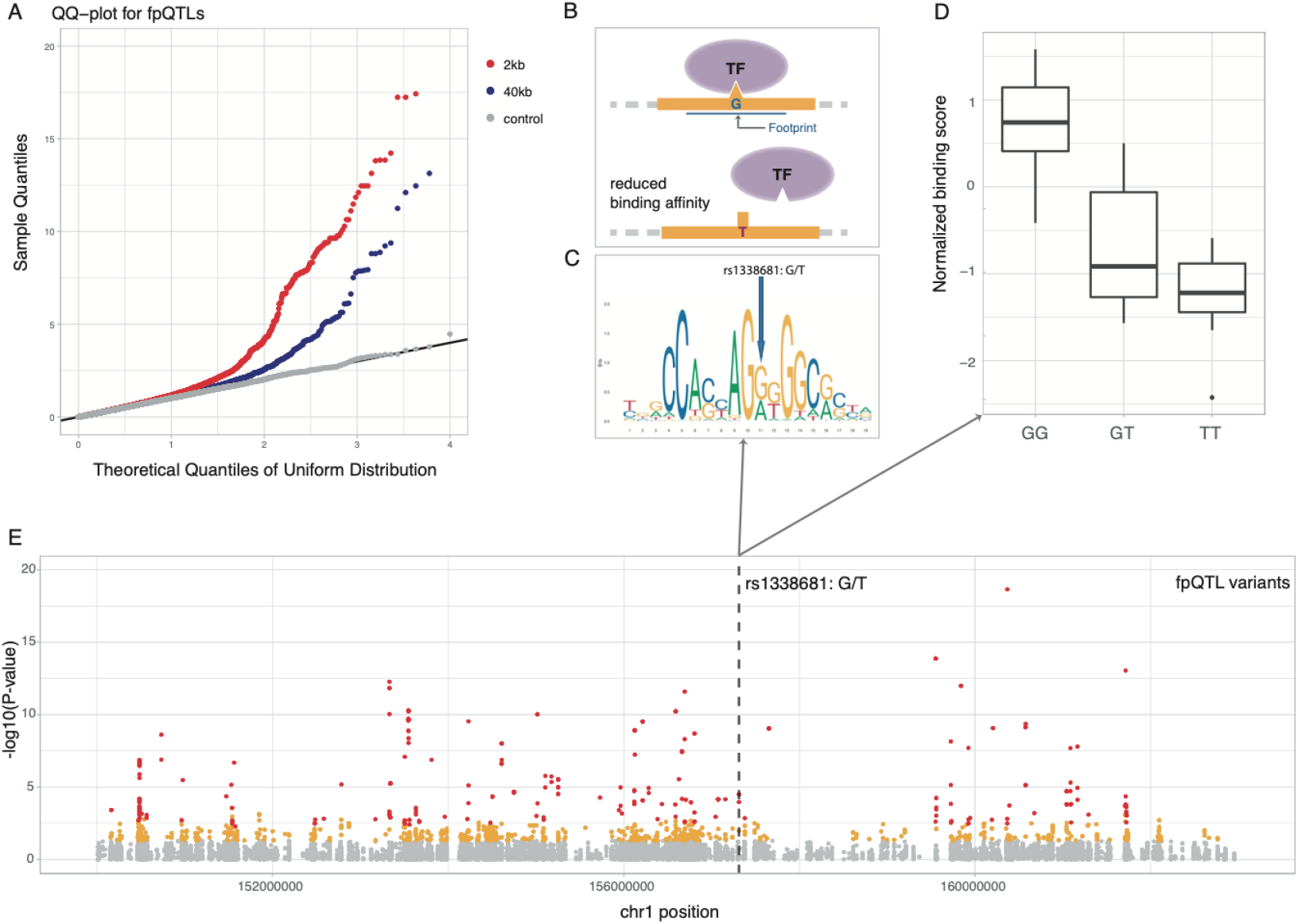
Genome-wide detection of fpQTLs and an example fpQTL SNP. (A) QQ-plots for all tests of association between footprint scores and variants within 2kb (red) and 40kb (blue) regions surrounding the target footprint. Permutation controls (gray) confirmed that observed p-values are uniform under the null (B, C, D) Example of a fpQTL rs1338681: (B) The T allele reduced TF binding affinity through (C) disrupting the CTCF binding motif, and (D)is associated with reduced binding score. (E) -log10 p values of associations of variants-footprint tested, significant fpQTLs at 10% FDR are in red color.

We first analyzed the genomes of 57 Yoruban individuals from HapMap, using published DNase-seq data from lymphoblastoid cell lines (LCLs) and genome-wide variant annotations from the 1000 Genomes Project. Annotations were prepared using the TRACE preprocessing pipeline modified to incorporate genome-wide variant data from all 57 individuals (see methods, **Supplemental Fig. 1**). Processed signals were subsequently used to predict individual-specific TFBS footprints with TRACE. Individual-specific marginal posterior probabilities for each region being bound by a certain TF wer also assigned to each corresponding predicted footprint.

For each TFBS within a sample, the TRACE score was used as a quantitative trait to estimate sample-specific TF binding activity at individual footprints. Given genome-wide footprint predictions and genotypes, association tests were performed between TRACE scores for each footprint-SNP pair in a *cis* candidate window of 2kb and 40kb centered at the target footprint (**Figure 1**). Each Footprint-SNP pair where variable TF binding affinity was significantly correlated with genome variants at 10% FDR were labeled as fpQTLs. In general, fpQTLs within a 2kb window showed higher significance levels than fpQTLs in a 40kb window (**Figure 1A**), therefore only fpQTLs within a 2kb window were retained for further analysis.

### Proximal fpQTLs have a larger effect on binding

The distance between a fpQTL SNP and its target TFBS may impact its effect size on binding activity. Since substitutions within a motif can directly lead to TFBS creation or disruption, we hypothesized that fpQTL SNPs within target footprints could have greater effects on TF binding than those outside footprints, and that effect sizes would be inversely related to distance from footprints. To test for these effects, fpQTLs were separated into three groups: 5.15% of SNPs were locating inside associated footprints, 29.2% were outside of the target footprint but fell within a +/-100 bp window centered at the footprint, and the remaining SNPs were outside the +/- 100bp window but were within a +/-1 kb window (**Figure 2A**). Each footprint can be linked to multiple SNPs, however, 12.5% of fpQTL loci were significantly linked to SNPs that fell within the target footprint, and 55.4% had associated SNPs lying in a +/-100 bp window centered at the footprint.

**Figure 2:**
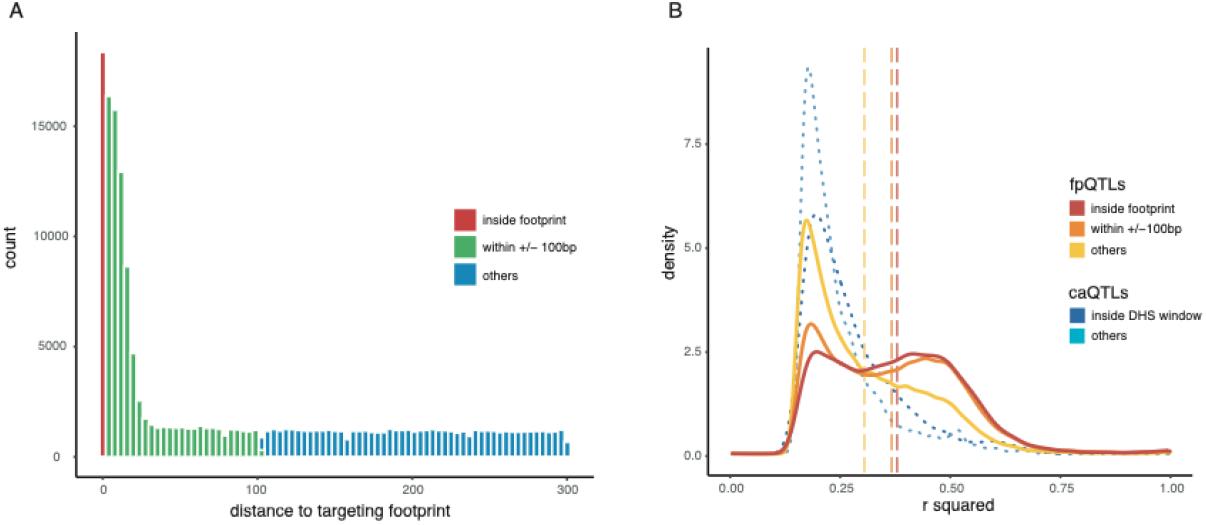
Properties of Proximal and distal fpQTLs. (A) Distribution of fpQTLs by variant distance to target footprint. (B) Distribution of effect sizes for proximal and distal fpQTLs (solid curves) and caQTLs (dotted curve) showed larger effect sizes for more proximal QTLs. Dashed lines are the average effect size for the three positional categories of fpQTLs.

To test if distance factors into the impact of genetic variants on footprints, we used the R^2^ from the linear model to indicate effect sizes. The distribution of effect sizes in three positional categories indicated that fpQTLs inside the binding sites tend to have a larger effect on binding activity, and more distal fpQTLs tend to have smaller effect size (**Figure 2B**). This is consistent with our expectations that formation or disruption of a binding sequence can exert the greatest effect on TF binding. The same trend had been observed with chromatin accessibility QTLs (caQTLs) as proximal SNPs have a more extreme effect on chromatin accessibility than more distant ones (**Figure 2B**, dotted curves) (Degner et al. 2012).

### Functional significance of fpQTLs

Identified fpQTLs provide a rich resource to generalize genomic properties of variants, including their enrichment with other cellular traits or diseases. fpQTLs share a large overlap with caQTLs that were also identified in LCLs using the same DNase-seq datasets. Utilizing GWAS catalog data, many fpQTL SNPs were found to overlap with GWAS-associated SNPs for various disease/traits including Type 2 diabetes and cancer such as prostate cancer, consistent with prior knowledge, as well as blood protein levels, mean corpuscular volume and serum metabolite levels. Fisher’s exact test also showed significant enrichment in some traits or diseases such as IgM levels and systemic lupus erythematosus (**Figure 3**).

**Figure 3:**
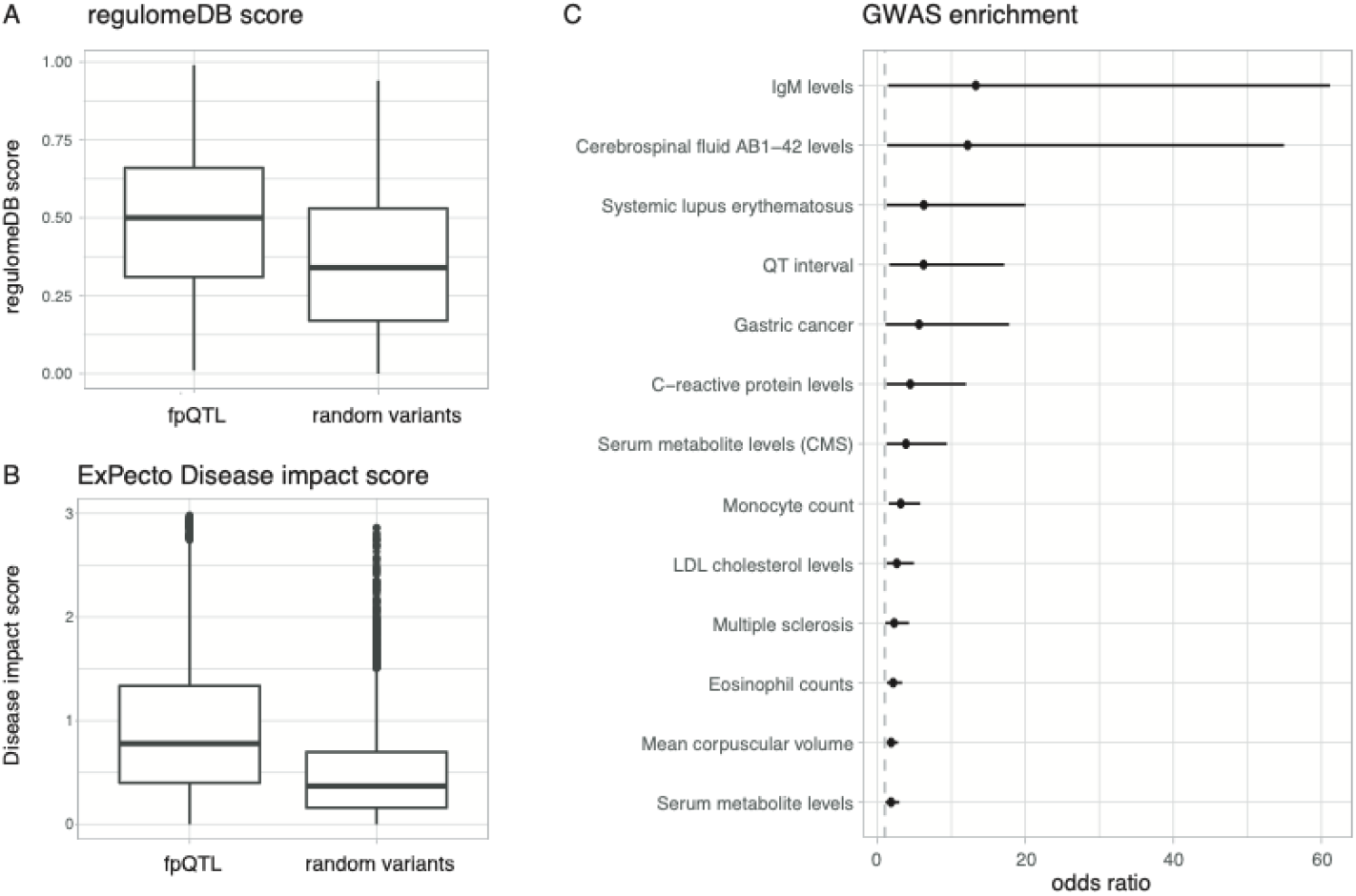
Functional significance of fpQTLs. (A) RegulomeDB scores of detected fpQTL SNP and random selected variants. (B) ExPecto disease impact score of detected fpQTL SNP and random selected variants. (C) GWAS catalog Disease or traits enrichment in fpQTLs from Fisher exact test (P value <0.02 and odds ratio > 1). Odds ratio and its 95% confidence interval are shown as dots and horizontal lines.

Several computational tools have been developed to prioritize genetic variants and predict their functional consequences based on a variety of genomic features. To study genomic properties of fpQTLs, fpQTL SNPs were queried against RegulomeDB (Boyle et al. 2012; Dong and Boyle 2022) and ExPecto annotations (Zhou et al. 2018), and functional scores were obtained for each SNP. Compared to randomly selected non-fpQTL variants, fpQTL SNPs showed higher functional significance in both scoring systems (**Figure 3**).

### fpQTL-eQTL pairs reveal positive and negative regulatory activity

Genetic variations that alter the likelihood of TF binding can, in turn, affect regulatory networks. Thus, we hypothesized that fpQTLs will also contribute to variation in the expression level of nearby genes. To examine the effect of fpQTLs as a potential mechanism linking noncoding variation with phenotypic variation, tissue specific eQTLs from Epstein-Barr virus (EBV) transformed lymphocytes were retrieved from the Genotype-Tissue Expression (GTEx) repository. We found that 12.8% of fpQTL SNPs are also significantly associated with variation in the expression levels of nearby genes. These fpQTL SNPs are enriched within 1KB of TSSs (9.2% are within +/- 1kb of the TSS; 78.6% are > 1kb distant, but within +/- 100kb), and strength of association was inversely correlated with distance from the TSS (**Supplemental Fig. 2**).

However, unlike dsQTL-eQTLs pairs, where increased chromatin accessibility is typically associated with increased gene expression, the direction of the fpQTL-eQTL SNP’s impact on gene expression is highly diverse. This likely reflects the indirect impact of fpQTL SNPs on gene expression, which are mediated by the specific activity of the TFs that binds their respective target footprints. Since TFs are highly diverse in their activating and/or repressive effects, increased binding affinity does not always translate to increased gene expression. Therefore, to study the correlation between the impact of SNPs on TF occupancy and gene expression levels, fpQTLs and eQTLs regression slopes were calculated from linear models of footprint scores and gene expression respectively. This method is not restricted to changes in a single direction, since the sign of the slope indicates either up-regulation or down-regulation of binding affinity or gene expression. Congruent signs for both slopes suggest a direct correlation between altered TF affinity at the target footprint for a fpQTL and target gene expression. For example, reduced binding affinity linked to reduced gene expression (**Supplemental Fig. 3A-C**). Conflicting signs, on the other hand, indicate inverse correlation between TF affinity and gene expression. For example, reduced TF binding activity associated with increased gene expression (**Supplemental Fig. 3D-F**).

A TF has directional impact on gene expression if changes in binding activity have a consistently positive or negative effect on gene expression across all matched fpQTLs. To identify such TFs, we performed a binomial test on each TF separately to detect those for which the signs of the TF-binding and transcript-abundance slopes were systematically more congruent or conflicting compared to a random expectation. There were 43 TFs with Bonferroni-corrected p-values smaller than 0.05 and a fold change greater than 2 (**Figure 4A**). Among these TFs, nuclear transcription factor Y, beta subunit (NFYB) has the most significant negative effect TF but has a moderate fold change of positive loci / negative loci. This suggests that, although NFYB loci are more enriched for having a negative regulatory effect, some still exhibited up-regulation on gene expression. In fact, the NF-Y complex is known as an activator protein that can bind promoter regions and regulate transcription through heterodimer or heterotrimer formation. However, the NFYB subunit of the NF-Y complex has been previously reported to be a repressor in multi-omics analysis (Zhao et al. 2016; Tharyan et al. 2020). Another significant directional TF is eomesodermin (EOMES) that act as a transcriptional activator and is involved in differentiation of CD8+ T cells (Atreya et al. 2007; Baala et al. 2007; Shimizu et al. 2019), consistent with its high positive log fold change value.

**Figure 4:**
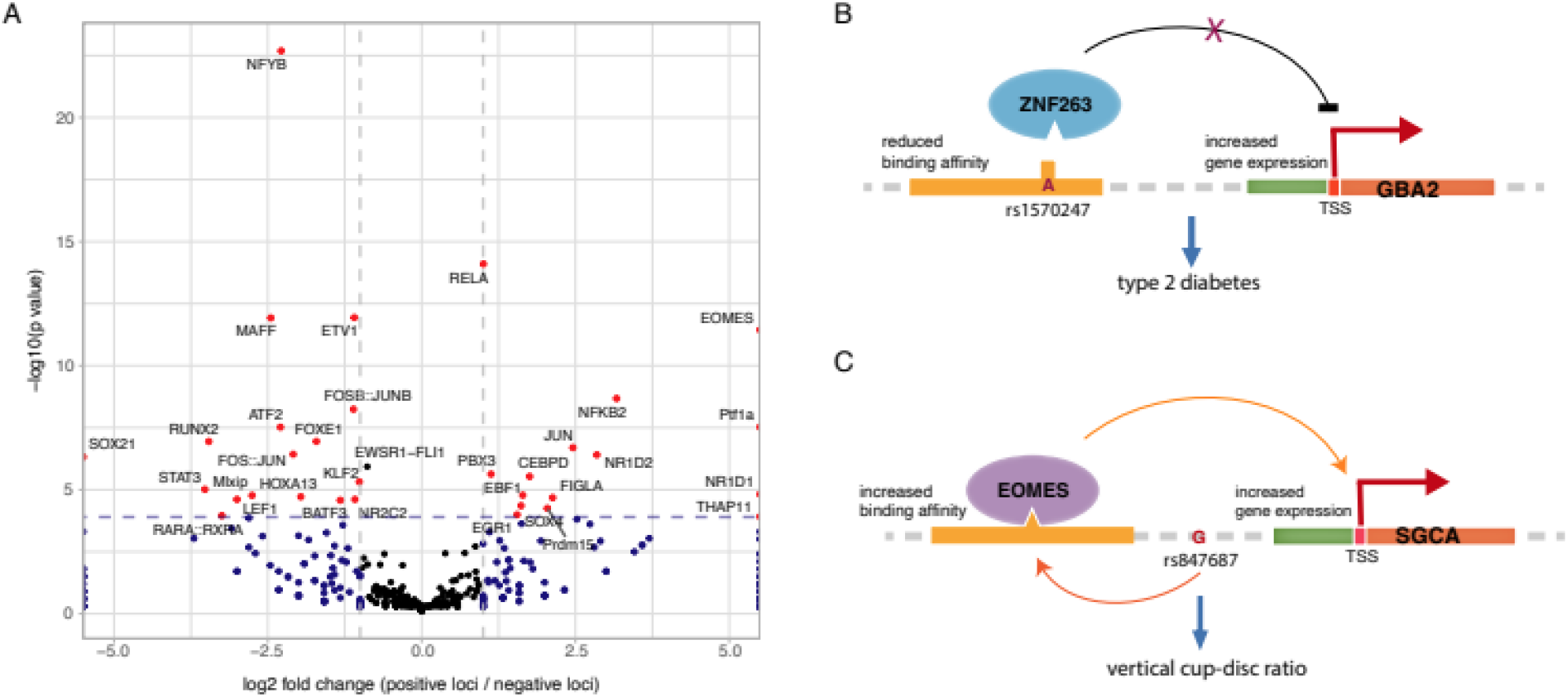
Properties of fpQTL-eQTL. (A) Volcano plot for TFs on their gene regulation directionality. (B) rs1570247 disrupts ZNF263 binding and is correlated with increased expression of GBA2 gene. This SNP is also linked to type 2 diabetes. (C) rs847687 is associated with increased binding of EOMES and increased expression of the SGCA gene. This SNP is linked to the vertical cup-disc ratio trait.

## Discussion

GWAS studies using sequencing-based assays such as RNA-seq, RIBO-seq, ChIP-seq and DNaseI-seq have revealed large numbers of QTLs involved in gene expression (Pickrell et al. 2010) and transcriptome (Montgomery et al. 2010; Lappalainen et al. 2013; Battle et al. 2014), ribosome occupancy and protein abundance (Battle et al. 2015), chromatin accessibility (Degner et al. 2012), histone modification (McVicker et al. 2013), and DNA methylation (Banovich et al. 2014). Here we define footprint QTLs (fpQTLs) as genetic variants that are significantly associated with TF footprint binding affinity, providing additional information on the regulatory impacts of genetic variation. Both proximal and distal fpQTLs were identified using association tests and, although proximal fpQTL SNPs are more-enriched and have greater effect sizes relative to their target fpQTL, we also detected many distal fpQTLs outside the 2kb window surrounding each fpQTL.

fpQTLs from this study provide a rich source of information for examination of SNP genomic properties and evaluation of the regulatory potential of untested variants to achieve a better understanding of regulatory mechanisms. Among fpQTL SNPs, many also overlapped with eQTL SNPs and potential disease-causing SNPs identified by GWAS. For example, rs1570247, a causal SNP correlated with type 2 diabetes, disrupts the binding of the previously reported transcriptional repressor ZNF263 (Weiss et al. 2020), and is associated with increased expression of the Glucosylceramidase Beta 2 (GBA2) gene (**Figure 4**). This provides a potential underlying mechanism for the linkage between genetic variants, gene expression and disease phenotype, as the alternative allele can alter ZNF263 binding. Subsequent disruption of ZNF263 binding can affect GBA2 gene expression along with other co-factors and cause the disease phenotype. Another example is rs847687, which increases the binding of the activating TF EOMES that was discussed in the previous section and is associated with increased expression of the Sarcoglycan Alpha (SGCA) gene. This has also been identified as a disease risk locus and was linked to vertical cup-disc ratio through the same gene by a GWAS study (Craig et al. 2020), postulating that EOMES binding might be involved in the cup-disc ratio phenotype via transcriptional regulation.

Until now, our ability to identify variants that affect TF binding was limited to variants within TFBSs themselves. TRACE scores change this by providing a quantitative phenotype that can be used to directly predict TF affinity for any given footprint. The fpQTLs derived from association tests using these scores represent a novel vantage point on how regulatory variants elicit their phenotypic effects. This method can reveal clues about the intricacies of how individual TFs function within a broader regulatory context, including the identity of potential cofactors and binding partners, associated histone modifications, chromatin properties, etc. We can use these clues to begin untangling the mechanisms involved in modulating the regulatory effects of different TFs. Using these insights, we can build a more-complete understanding of the genetic regulatory code by which transcriptional outcomes are determined.

## Methods

### Data and software

DNase-seq data for 57 HapMap Yoruban lymphoblastoid cell lines (LCLs) in fastq format were retrieved from http://eqtl.uchicago.edu/dsQTL_data/. Corresponding variant files in VCF format were downloaded from 1000 Genomes. Reads were mapped to hg38 using the variant-aware aligner VG toolkit (Garrison et al. 2018). 780 PWMs and cluster information (Supplemental Table S2) were downloaded from the JASPAR database (Khan et al. 2018).

### Modified TRACE workflow

Signal processing of DNase-seq data followed the TRACE pipeline, which includes cutting bias correction, normalization, and local regression smoothing. The original program was modified to accept sequence information with variants, and the motif score feature can reflect different alleles. To generate individual-specific TFBSs, a 10-motif was trained for each TF and the same model was applied on all individual datasets.

### fpQTL association testing

For each predicted footprint and its individual-specific binding score from TRACE, we tested for association of the binding score with the genotypes of all SNPs where the minor allele frequency was greater than 5% within a cis-candidate region of 2kb or 40kb, centered on the target footprint. For each footprint and each SNP falling within the candidate window, standard linear regression between genotype and binding score was performed in R, and a p-value was generated by testing the alternative hypothesis that the slope in the linear regression model is not 0. The “qvalue” R package was used to estimate a significance threshold for footprint-variant pairs corresponding to a 10% FDR.

### caQTLs generation

To generate caQTLs, DNase-seq reads that were mapped to the hg38 assembly in the previous section were processed following the same pipeline described in (Degner et al. 2012). Reads starting within the 5 bp window centered at SNPs were discarded. Raw DNase sensitivity was calculated by counting the number of reads falling in each non-overlapping 100bp window, normalized by total number of mapped reads and mappability. Additional normalization steps for hypersensitivity phenotypes include GC-content correction, mean-center and variances scale, and quantile-normalization to standard normal distribution. Unidentified confounders were removed with PCA, thus 4 PCs were removed. Subsequent association testing was conducted to generate caQTLs at 10% FDR.

### Functional significance analysis for fpQTLs

Genomic annotations and functional scores were collected from RegulomeDB (Boyle et al. 2012; Dong and Boyle 2022) and ExPecto (Zhou et al. 2018). All identified fpQTL SNPs and randomly selected background non-fpQTL SNPs were queried, and lists of RegulomeDB scores and ExPecto disease significance scores were obtained. GWAS Catalog data (Zhou et al. 2018; Welter et al. 2014) for the Dec 2021 build were retrieved from the GWAS Catalog website. Fisher’s exact tests were performed to assess the enrichment of disease and other traits. Odds ratio and its 95% confidence interval were calculated for each disease/trait.

### Impact direction of the fpQTLs-eQTLs SNPs

The impact direction of the fpQTLs-eQTLs SNPs is defined as the consistency of the signs of the slopes of the variant-footprint and variant-expression linear regression model. It can be interpreted as the up or down-regulation effect of the increased or reduced TF binding resulting from the alternative allele relative to the reference allele on gene expression (i.e., if it has a negative impact direction, the disruption of TF binding will lead to increased expression of a nearby gene). Log fold-change for each TF was calculated by taking the log-ratio of the number of fpQTLs-eQTLs SNPs associated with that specific TF that have a positive impact direction to the number of SNPs with negative effects.

## Data and code availability

TRACE is an open source software; the source code, trained models, and predictions are available on GitHub (https://github.com/Boyle-Lab/TRACE).

## Acknowledgments

We thank Shengcheng Dong for her help in generating functional annotation scores. Special thanks is extended to Adam G. Diehl for editorial support and feedback on the manuscript. We also want to thank all members of the Boyle Lab for their support and constructive feedback.

This project was supported by NIH U24 HG009293 and the Eleanor and Larry Jackier U-M/Technion and Weizmann Collaborative Research Grant.

