## Supplemental Materials for "Quantitative assessment of association between noncoding variants and transcription factor binding"

for

### Supplemental Figures

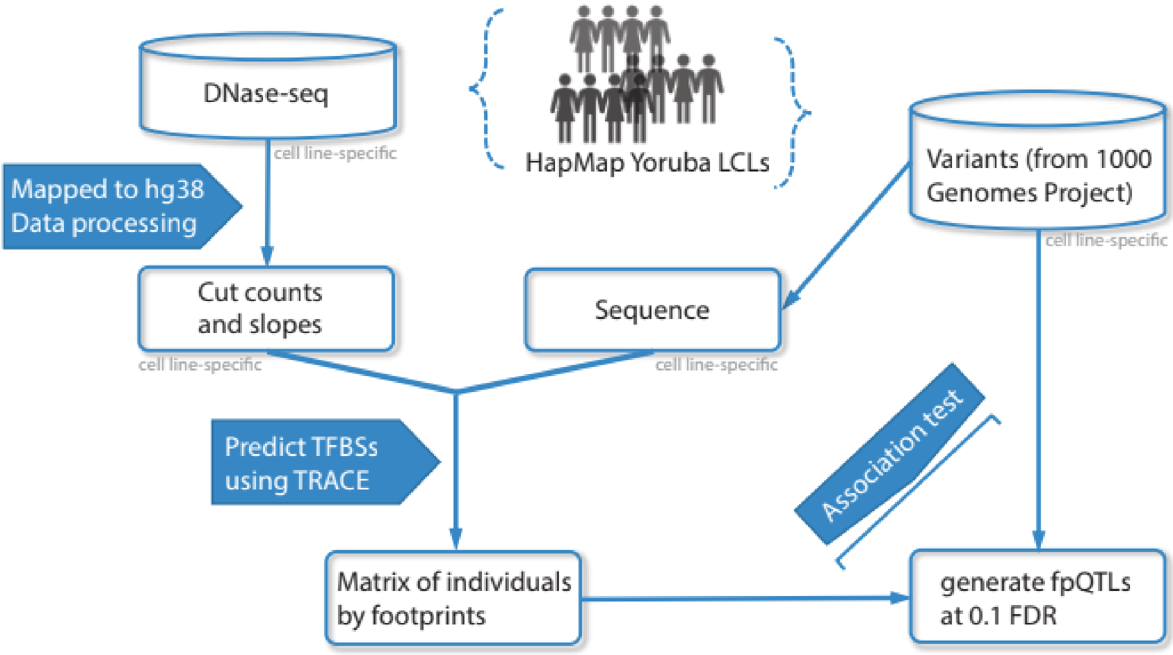


**Suppplementary Figure 1: workflow of fpQTLs identification**

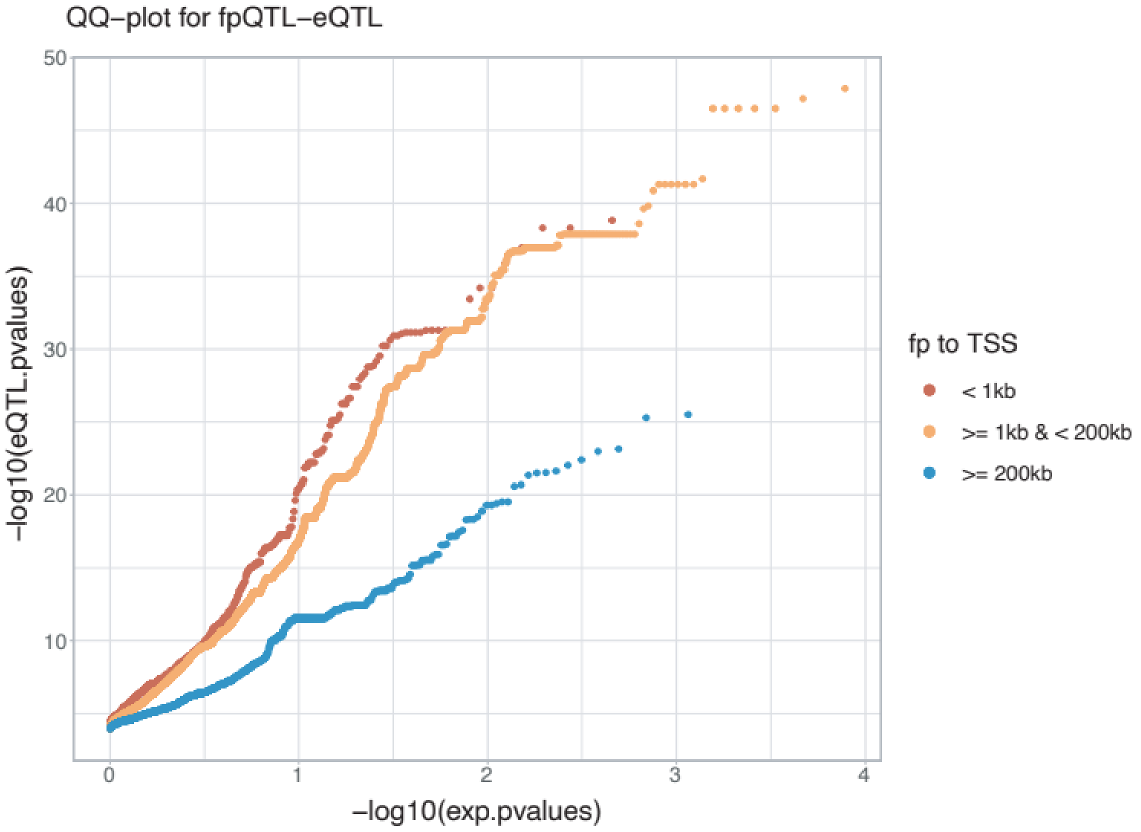


**Supplementary Figure 2: QQ-plots for fpQTL-eQTL pairs with significant p values, by distance to the nearest TSS.**

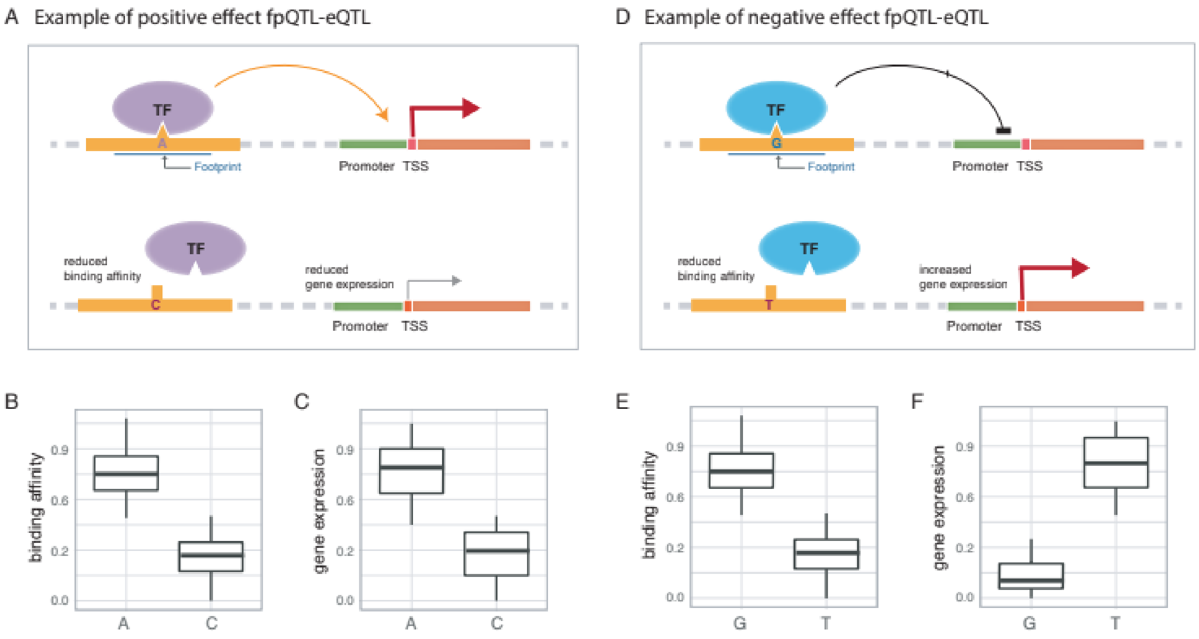


**Supplementary Figure 3: Simplified schematic of (A, B, C) positive and (D, E, F) negative impact fpQTL-eQTL.** (A) a positively directed genetic variant disrupts TF binding at footprint and decreases expression of the gene regulated by that regulatory element. the A/C allele is associated with (B) level of TF binding and (C) gene expression. (A) a negatively directed genetic variant disrupts TF binding at footprint and induced gene expression. the G/T allele is associated with (B) level of TF binding and (C) gene expression.
